# Human acute microelectrode array recordings with broad cortical access, single-unit resolution and parallel behavioral monitoring

**DOI:** 10.1101/2022.10.27.514100

**Authors:** Viktor M. Eisenkolb, Lisa M. Held, Alexander Utzschmid, Xiao-Xiong Lin, Sandro M. Krieg, Bernhard Meyer, Jens Gempt, Simon N. Jacob

**Author notes:** These authors contributed equally to this work (Jens Gempt, Simon N. Jacob).

## Abstract

Human single-unit studies currently rely on neurosurgical procedures that provide only limited brain coverage and on recording devices that do not integrate easily into established surgical routines. Here, we report reliable and robust acute multi-channel recordings with broad cortical access using planar microelectrode arrays (MEA) implanted intracortically in awake brain surgery. We provide a comprehensive characterization of extracellular neuronal activity acquired intraoperatively in tumor patients with large open craniotomies. MEA implantation was fast, safe and yielded high-quality signals at the microcircuit, local field potential level, and at the cellular, single-unit level. Recording from parietal association cortex, a region previously unexplored in human single-unit studies, we demonstrate applications on these complementary spatial scales and describe travelling waves of oscillatory activity as well as single-neuron and neuronal population responses during numerical cognition including operations with uniquely human number symbols. Intraoperative MEA recordings are practicable and can be scaled up to explore cellular and microcircuit mechanisms of a wide range of human brain functions.

## Introduction

There are vast gaps in our understanding of the organization and operation of the human nervous system at the level of individual neurons and their networks. Limited opportunities to directly access the human brain call for multidisciplinary collaborations that combine expertise in neuroscience and clinical medicine to invasively measure neuronal activity with single-unit resolution (Cash & Hochberg, 2015). This approach has been most fruitful in patients with medically intractable epilepsy implanted with microwire bundles (Fu et al., 2022; Kaminski et al., 2017; Kornblith et al., 2017; Kutter et al., 2018; Minxha et al., 2020; Rutishauser et al., 2010; Sheth et al., 2012) and in patients with movement disorders undergoing deep brain stimulation (DBS) (Jamali et al., 2019; Jamali et al., 2021; Zaghloul et al., 2009). Two crucial challenges persist, however, in the investigation of the cellular and circuit physiology of human brain functions. First, epilepsy and DBS surgeries do not provide comprehensive brain coverage, leading to strong focusing of current human single-unit studies on the medial temporal lobe (MTL) and on small circumscribed regions of the frontal lobe. Second, reliable and robust recording technology is still lacking, meaning that clinicians must be trained on increasingly complex devices that necessitate significant modifications to standardized and proven surgical procedures (Chung et al., 2022; Paulk et al., 2022).

Broad access to the human cortex in large patient groups combined with easy-to-implement methods would greatly accelerate progress in researching the neuronal basis of human brain functions. Here, we demonstrate acute recordings from planar multi-channel microelectrode arrays (Utah MEAs) implanted intracortically in patients operated awake for the removal of left-hemispheric brain tumors. Tumor surgeries with open craniotomies expose large areas of cortex and allow for flexible placement of recording devices, meaning that electrode positions can be adapted to research questions - not vice versa. Awake surgeries with intraoperative functional mapping minimize the risk of postoperative deficits by delineating functionally important regions and thus increase the precision of tumor resection (Sanai et al., 2008). Patients undergoing awake surgery can perform a wide variety of tasks tapping into sensorimotor functions, visuospatial functions, language and other higher cognitive functions (Mandonnet & Herbet, 2021). Penetrating, intracortical MEAs are widely used for chronic measurements of single-unit and population activity in non-human primates (Chen et al., 2020; Mitz et al., 2017) and have shown potential for clinical applications (Schevon et al., 2019; Truccolo et al., 2011) as well as for neurorestorative brain-computer-interfaces (BCIs) in humans (Aflalo et al., 2015; Fernandez et al., 2021; Flesher et al., 2016; Hochberg et al., 2006; Pandarinath et al., 2017; Willett et al., 2021).

Despite these successes, acute intraoperative MEA recordings to investigate human brain functions have not been reported. Cortical microtrauma and neuronal ‘stunning’ are believed to prohibit measurements with these devices shortly after implantation (Fernandez et al., 2014; House et al., 2006). In this study, we show that these obstacles can be overcome with appropriate choice of the arrays’ geometrical configuration. All implanted arrays recorded high-quality extracellular signals at the microcircuit level (local field potentials, LFPs). MEAs with increased electrode spacing, however, outperformed standard arrays with higher densities and also captured activity at the cellular, single-unit level. To demonstrate applications on these complementary spatial scales, we describe oscillatory dynamics in the form of waves of activity travelling across human parietal association cortex, a region previously unexplored in human single-unit studies, and investigate single-neuron mechanisms of numerical cognition including operations with uniquely human symbolic quantities. Our findings demonstrate that intraoperative MEA recording technology is suited to provide the high-volume recordings necessary to advance translational research on the cellular and microcircuit basis of a wide range of human brain functions.

## Results

### Intraoperative MEA implantation

Awake surgeries with open craniotomies enable direct, controlled investigations of human brain functions while the patients are alert and can perform tasks of varying complexity (Mandonnet & Herbet, 2021) (Fig. 1A). Craniotomies overlap in particular over the motor cortical regions and over the posterior frontal lobes (Fig. 1B). They can extend anteriorly to the frontal pole and posteriorly to the parieto-occipital junction, dorsally to the inter-hemispheric fissure (midline) and ventrally to the temporal lobe. Typical craniotomies expose large regions of cortex (several tens of cm^2^), yielding broad access to the human brain. Infrared thermal imaging during a representative surgery verified that physiological temperatures are maintained at the cortical surface (Fig. 1C).

**Fig. 1.**
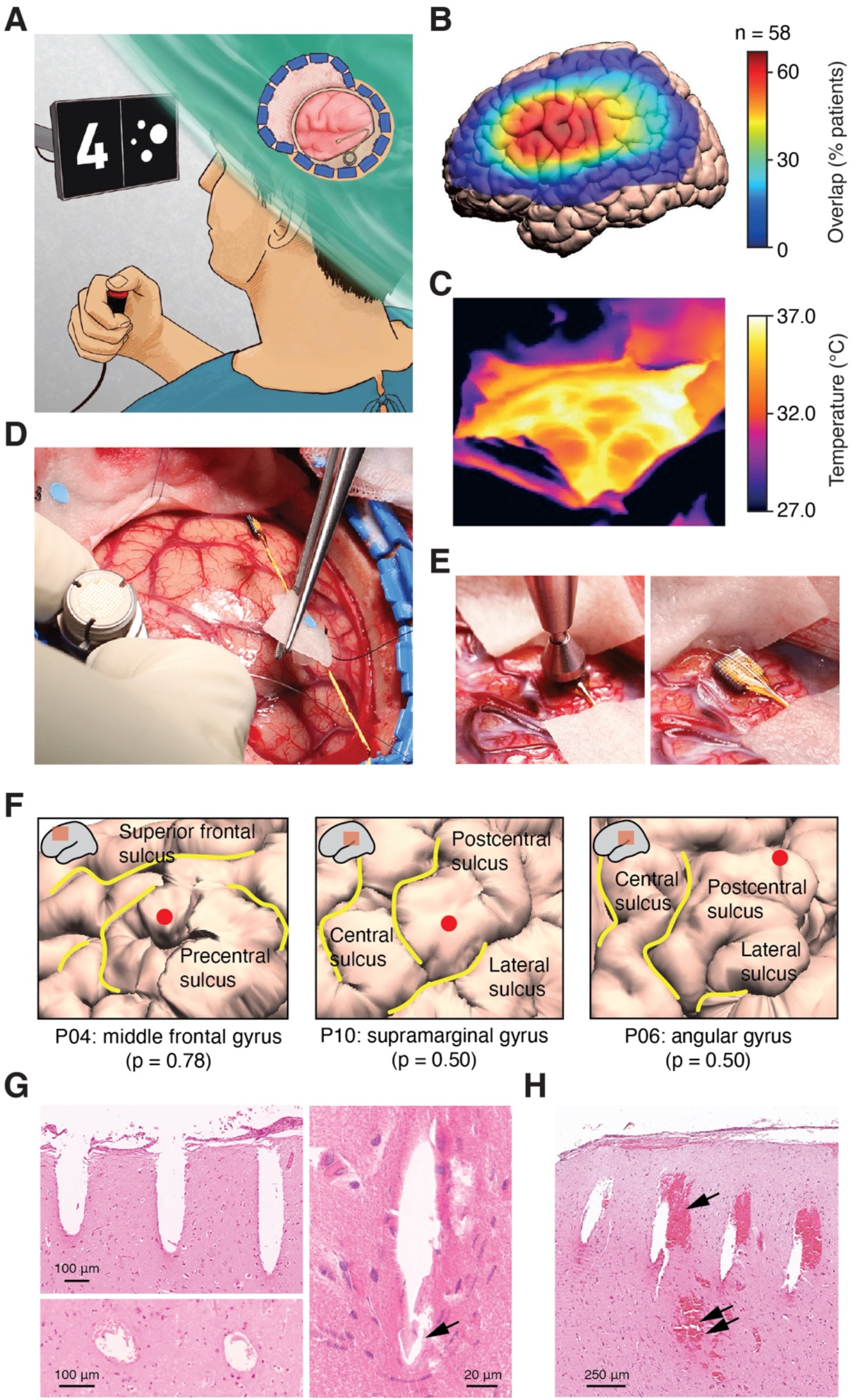
Awake brain surgery and intraoperative microelectrode array implantation. (**A**) Schematic of awake brain surgery providing access to the human cortex for microelectrode recordings in participants who can perform cognitive tasks. (**B**) Overlap of craniotomy locations in neurosurgical patients operated awake for the removal of left-hemispheric brain tumors (n = 58 surgeries performed in our department over the course of five years) projected onto the ICBM template brain. (**C**) Infrared thermal imaging of the cortical surface during a typical craniotomy procedure. (**D**) Placement of the microelectrode array in preparation of implantation. (**E**) Pneumatic insertion of the microelectrode array into cortex. (**F**) Cortical surface reconstruction of the implantation site in three example participants. The probability of implantation in the specified gyrus is given according to the JuBrain probabilistic cytoarchitectonic map. (**G**) Histological sections of an example implantation site showing electrode tracts as they penetrate the pia mater (top left, longitudinal section), along the electrode shaft (bottom left, axial section) and at the electrode tip (right, arrow). (**H**) Histological section of a different implantation site showing microhemorrhages along the electrode tracts (single arrow) and in deeper cortical layers (double arrow).

We performed a total of 13 acute microelectrode array (MEA) implantations in patients undergoing surgery for brain tumor resection, eight of which were operated awake (Table 1). Except for the procedures related to the array implantation, the course of the surgery was not changed. Following skin incision, preparation and opening of the skull and dura mater, but before awakening the patient from anesthesia, we placed the array’s pedestal next to the craniotomy, anchored it with skull screws and positioned the MEA over the target cortical area (Fig. 1D). Reference wires were inserted under the dura. We intended for the implantation site to lie as remotely as possible from the bulk tumor tissue but still within the pre-operatively determined resection area. The array was then pneumatically inserted and covered with saline irrigated strips (Fig. 1E) until explantation, typically when tumor resection started. With established and practiced procedures, the implantation could be performed in less than ten minutes. We encountered no adverse clinical events in connection to MEA implantation or recordings, neither during the surgery nor during routine patient follow-up over several months.

**Table 1.** Study participants.

| ID | Sex | Age | Tumor location | Procedure | State | Array location | Channels | Spikes | Behavior | Notes |
| --- | --- | --- | --- | --- | --- | --- | --- | --- | --- | --- |
| P01 | F | 68 | right frontal | histology | anesthetized | inferior parietal cortex | 96 | N/A | N/A |  |
| P02 | M | 54 | right parietal | histology | anesthetized | inferior parietal cortex | 96 | N/A | N/A |  |
| P03 | M | 62 | right parietal | histology | anesthetized | inferior parietal cortex | 96 | N/A | N/A |  |
| P04 | M | 56 | left frontal | setup testing and recording | anesthetized | middle frontal gyrus | 96 | no | N/A |  |
| P05 | F | 75 | left central | setup testing and recording | anesthetized | superior frontal gyrus | 96 | no | N/A |  |
| P06 | M | 57 | left parietal | recording | awake | angular/supramarginal gyrus | 96 | (yes) | number task | spiking activity on very few channels only |
| P07 | M | 73 | left parietal | recording | awake | angular /supramarginal gyrus | 96 | no | number task | performance non-symbolic trials ↓ |
| P08 | F | 55 | left parietal | recording | awake | inferior parietal cortex | 96 | N/A | N/A | no data acquisition<br>bad ground |
| P09 | M | 51 | left fronto-parietal | recording | awake | middle frontal gyrus | 96 | no | number task | performance non-symbolic trials ↓ |
| P10 | M | 32 | left temporal | recording | awake | supramarginal/angular gyrus | 25 | yes | number task |  |
| P11 | M | 67 | left frontal | recording | awake | supramarginal/angular gyrus | 25 | yes | number task |  |
| P12 | M | 71 | left insular | recording | awake | angular/supramarginal gyrus | 25 | N/A | N/A | no data acquisition<br>intracerebral hemorrhage<br>(unrelated to implantation) |
| P13 | F | 59 | left central | recording | awake | supramarginal/postcentral gyrus | 25 | yes | number task | sudden SNR drop |

For each participant, the implantation site was reconstructed using intraoperative photographic documentation as well as pre-operative structural MR imaging. Three implantations were located in frontal cortex and ten in parietal cortex (Table 1). Examples of implantations in the middle frontal gyrus, the supramarginal gyrus and the angular gyrus are shown (Fig. 1F).

We histologically analyzed three implantations (Table 1). Grids of electrode tracts could be clearly identified from the penetration of the pia mater along the course of the shafts to - in some instances - the tip of the electrode (Fig. 1G). In two patients, cortical tissue surrounding the electrodes showed no structural abnormalities across the entire array. In one patient, we observed petechial micro-hemorrhages along several electrode tracts as well as in deep cortical layers (Fernandez et al., 2014; House et al., 2006) (Fig. 1H). However, these changes were strictly confined to the vicinity of the electrodes. We did not detect any pathology distant from the implantation site.

In sum, implantation of intracortical MEAs in patients undergoing awake brain surgery is safe and practicable, achieving broad and direct access to the neuronal networks of the human cortical left hemisphere.

### Extracellular signal quality on MEAs with differing geometrical configurations

In the group of patients operated for awake tumor resection, we discontinued the anesthesia following MEA implantation. We began recording wide-band extracellular activity (Fig. 2A) as soon as the patients were alert and able to engage in conversation with the clinical team and prior to cortical electrostimulation for mapping of language-associated areas. Typically, the arrays had been settling for 30 to 40 minutes. We emphasize that the surgery was not prolonged by this time period; we merely used the awakening time to allow for the signals to develop and stabilize.

**Fig. 2.**
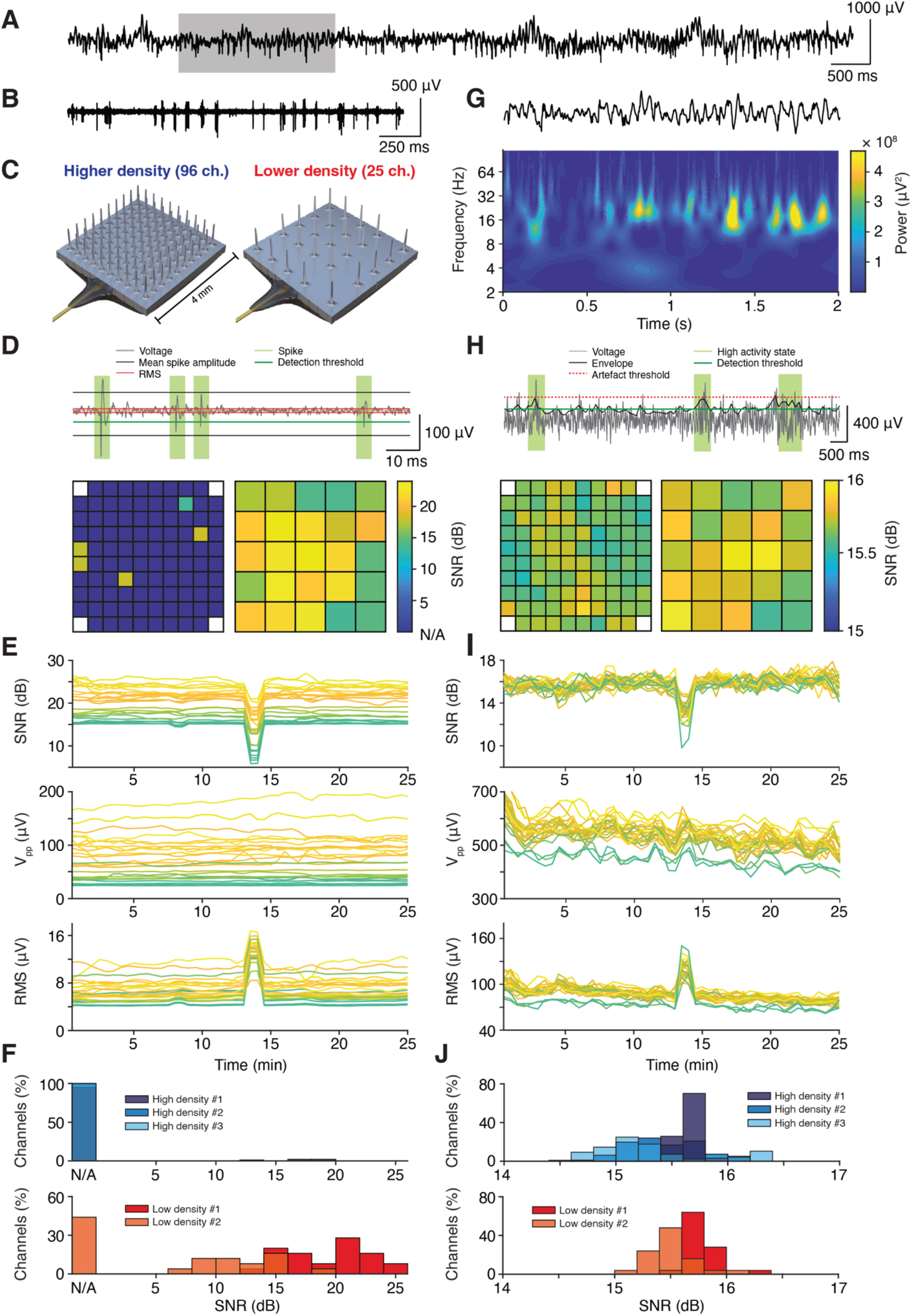
Extracellular neuronal signals recorded from microelectrode arrays with different densities. (**A**) Wide-band extracellular voltage signal recorded at an individual electrode (10 s trace). (**B**) High-pass filtered signal showing extracellular spiking activity in the section highlighted in (A) (2 s trace). (**C**) CAD drawings of the standard higher-density microelectrode array (left, 96 active channels) and of the custom lower-density microelectrode array (right, 25 active channels) used for intraoperative recordings. (**D**) Top: Schematic of the procedure for identifying spikes in high-pass filtered voltage signals. Bottom: Session-averaged SNR of a representative higher-density and a lower-density array (left and right, respectively). (**E**) Time course of spike SNR (top), peak-to-peak amplitude (middle) and RMS noise (bottom) across the entire session recorded with the lower-density array in (D). Note the brief increase in noise and reduction in SNR in the middle of the recording. (**F**) Distribution of spike SNR values obtained from electrodes in higher-density and lower-density recordings (top and bottom, respectively). (**G**) Low-pass filtered signal showing oscillatory LFP activity in the section highlighted in (A) (2 s trace). (**H**) Top: Schematic of the procedure for quantifying SNR in low-pass filtered voltage signals. Bottom: Session-averaged SNR of a representative higher-density and a lower-density array (left and right, respectively; same arrays as in (D)). (**I**) Time course of LFP SNR (top), peak-to-peak amplitude in high activity states (middle) and RMS in low activity states (bottom) across the entire session recorded with the lower-density array in (D). Note the same deflections in LFP noise and SNR as in the spike-filtered signal in (E). (**J**) Distribution of LFP SNR values obtained from electrodes in higher-density and lower-density recordings (top and bottom, respectively).

We first sought to evaluate the ability to detect the activity of individual neurons (i.e. spikes), present in the high frequency signal components (high-pass filter 250 Hz; Fig. 2B-F). We compared two different MEA configurations: a standard, higher-density array with 400 μm electrode spacing (pitch) and 96 active channels on a 10×10 grid and a custom, lower-density array with 800 μm pitch and 25 channels (Fig. 2C left and right, respectively). We performed four implantations with each array type (Table 1). Technical difficulties with grounding (P08, higher-density array) and a medical complication not related to the implantation (P12, lower-density array) did not allow us to advance to neuronal recording in two surgeries. In one case, we observed an abrupt drop in signal quality a few minutes into data acquisition (P13, lower-density array), prompting us to omit this data set from in-depth analysis. Qualitatively, prior to the unexplained event, the recording was not different from the other lower-density recordings.

The likelihood of recording spiking activity varied significantly between array configurations. In an example higher-density array, spiking activity of sufficiently high amplitudes for subsequent waveform sorting was present in only a few channels (Fig. 2D, left). In contrast, in an example lower-density array, spikes were detected on all electrodes (Fig. 2D, right). SNRs in this array were stable across the entire recording (25 minutes), with the exception of a single large electrical artefact leading to an increase in noise (Fig. 2E; Fig. S1A, B). This did not impact spike amplitudes, however, which remained stable during data acquisition. Across all successful recordings, this pattern was reproduced (Fig. 2F): in three consecutive implantations with the higher-density array (five implantations including two anesthetized participants, Table 1), we did not observe appreciable spiking activity (2 % of channels). In three consecutive implantations with the lower-density array (one recording not shown due to early termination, see above), we obtained spikes on the majority of channels (78 % of channels; p < 0.001, Fisher’s exact test higher-density vs. lower-density arrays). In the event that spiking activity could be recorded, SNRs were comparable (mean 17.1 ± 0.9 dB and 16.8 ± 0.8 dB for higher-density and lower-density arrays, respectively; p = 0.91, two-tailed Wilcoxon test).

Next, we evaluated the quality of LFPs, a measure of local network activity, i.e. the low-frequency component of our extracellular recordings (low-pass filter 250 Hz; Fig. 2G-J). Epochs of increased LFP activity were readily detected in both higher-density and lower-density arrays and across all channels (Fig. 2H; same example arrays as in Fig. 2D). In both array configurations, SNRs were high and displayed spatial clusters of similar signal strength. In the lower-density array, the clusters of high spiking SNR and high LFP SNR overlapped. As for the spiking activity, LFP signals were stable across the recording session and affected only momentarily due to a single electrical artefact (Fig. 2I; Fig. S1A, B). Across all successful recordings, LFP SNRs were very uniform across channels (mean 15.5 ± 0.1 dB and 15.7 ± 0.03 dB for higher-density and lower-density arrays, respectively; Fig. 2J).

Overall, electrical artefacts could be well controlled during intraoperative data acquisition. Very rarely, we observed a single high-amplitude ‘pop’ across all electrodes that disrupted recordings for a few hundred milliseconds until the signals settled again (Fig. S1A, B). 50 Hz line noise and its harmonics were regularly present in the recordings (Fig. S1C, D), but could be efficiently removed by offline filtering. Good grounding (i.e. strong connection of the pedestal to the skull) significantly reduced the hum. Bad choice of grounding, in contrast, lead to signal contamination, e.g. by facial muscle activity (Fig. S1E, F).

To determine whether single units could be isolated from the population (multi-unit) spiking activity (Fig. 3A), we sorted the thresholded waveforms. Distinct waveform clusters representing well-isolated single units were separated from noise (Fig. 3B, C) with little to no loss of spikes around the detection threshold (false negatives, Fig. 3D; less than 5 % of spikes in 74 % of units), no contamination by spikes violating the refractory period (false positives, Fig. 3E; less than 1 % of spikes in all units), stable firing rates throughout the recording session (Fig. 3F) and little to no mixing of spikes between different clusters (Fig. 3G). Following this procedure, single units could be isolated on the majority of electrodes in the example lower-density array (Fig. 3H), with two or more single units present on multiple channels. Across all analyzed recordings, single units were rarely picked up by the higher-density arrays (2 % of channels) but frequently isolated on the lower-density arrays (62 % of channels; p < 0.001, Fisher’s exact test higher-density vs. lower-density arrays). On lower-density array electrodes with sortable spikes, we recorded on average 1.6 single units per electrode.

**Fig. 3.**
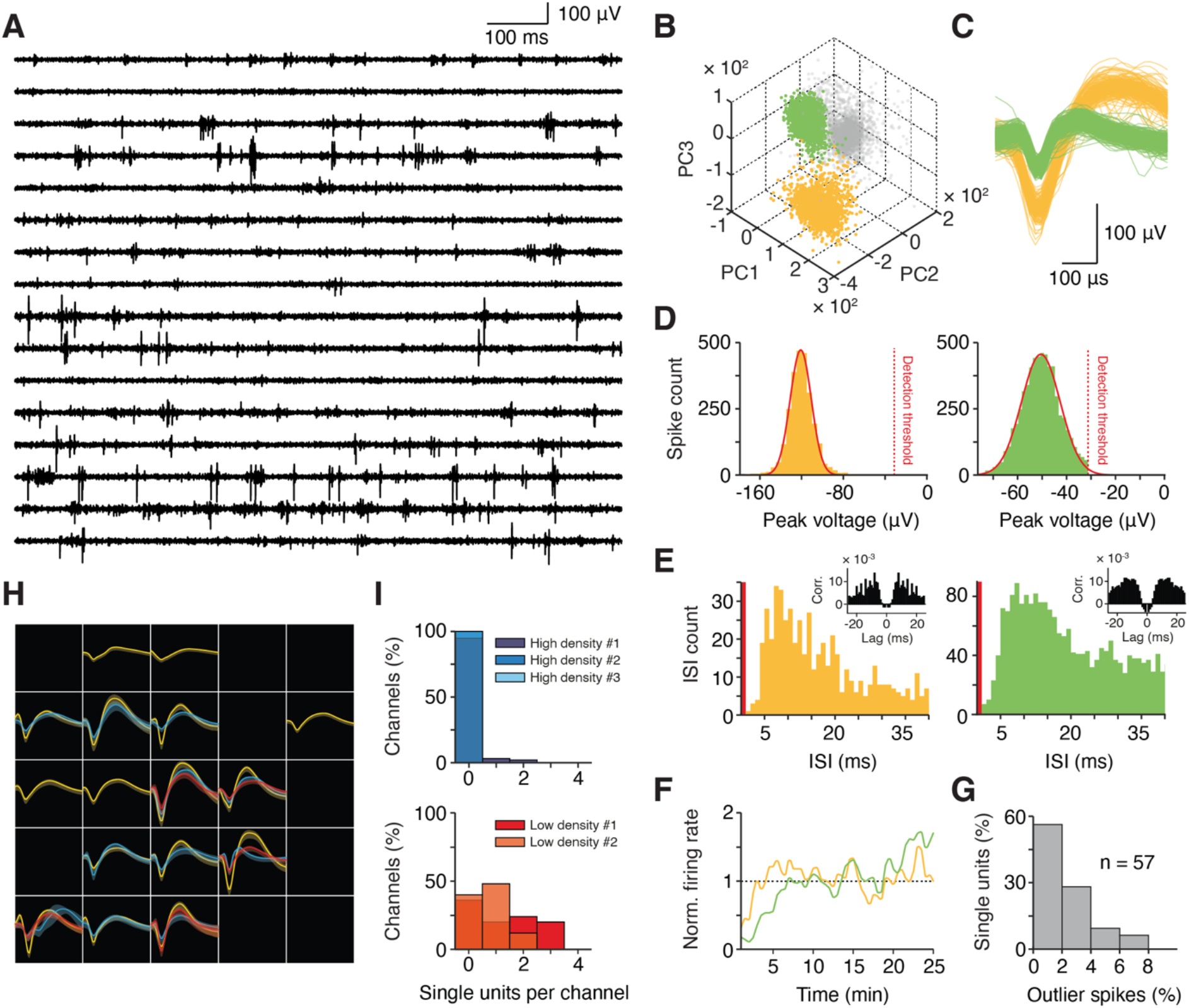
Isolation of single units from intraoperative microelectrode recordings. (**A**) High-pass filtered extracellular voltage signals from selected electrodes (1 s traces). (**B**) Principal component decomposition of thresholded waveforms recorded on an individual channel showing two distinct waveform clusters (yellow, green) separated from noise (gray). (**C**) Waveforms of the single units isolated by PCA in (B). (**D**) Distribution of waveform negative peak (trough) voltages for the two example units with gaussian fits and the selected detection threshold. (**E**) Distribution of inter-spike-intervals (ISI) for the two example units together with spike train autocorrelograms (insets). The refractory period (ISI < 1 ms) is marked in red. (**F**) Firing rates of the two example units across the entire recording session, normalized to a unit’s session-averaged activity. (**G**) Distribution of the percentage of spikes per unit that are assigned to different waveform clusters and thus considered outliers (n = 57 sorted units in all recordings). (**H**) Average single unit waveforms recorded from a lower-density microelectrode array. Bands indicate standard deviation across waveforms. Channels with multi-unit activity, but no well-isolated single units, are black. (**I**) Distribution of channels with well-isolated activity of one or more single units recorded from higher-density and lower-density arrays (top and bottom, respectively).

While single neurons represent the brain’s elementary processing units, it is increasingly recognized that temporal coordination and synchronization of neuronal activity across distances is crucial in particular for higher cognitive functions (Fries, 2015). Given their planar, grid-like configuration with well-defined spatial relationships between individual electrodes, MEAs are ideally suited to investigate the lateral propagation of activity in cortical networks. Several studies with chronic MEA recordings have reported waves of oscillatory brain activity that travel across the non-human primate and human cortex (Bhattacharya et al., 2022; Rubino et al., 2006; Sato et al., 2012; Takahashi et al., 2011) and could reflect higher-order organization of neuronal processing in space and time (Muller et al., 2018). Examination of oscillatory beta activity (20 ± 1.5 Hz) in a higher-density recording showed LFP peaks temporally shifted across neighboring electrodes with ordered progression of activity from the top to the bottom of the array (Fig. 4A). At each timepoint, LFP phases across the array could be approximated by a linear plane with non-zero slope aligned to the direction of activity propagation, in agreement with the notion of a travelling wave. We extracted and characterized such travelling waves in 500 ms epochs following presentation of visual stimuli (sample numbers, see Fig. 5) for both theta (6 - 9 Hz) and beta LFP bands (15 - 35 Hz; Fig. 4B-E). Waves travelled in preferred directions (p < 0.001 in theta and beta, Hodges-Ajne test for nonuniformity) that were frequency-band-specific (Fig. 4B). A second modal direction almost opposing the dominant primary direction suggested a spatial propagation axis (Fig. 4B), in line with intracranial EEG and ECoG recordings (Das et al., 2022; Zhang & Jacobs, 2015; Zhang et al., 2018) and during ictal discharges in patients with epileptic seizures (Liou et al., 2017; Smith et al., 2016). With increasing oscillatory frequency, travelling waves were detected less often (Fig. 4C) and showed higher propagation velocities (theta mean 0.57 m/s, beta mean 2.40 m/s; Fig. 4D), again matching data from chronic implantations. Spatial phase gradients fit the plane model well in both frequency bands (measured by Phase-Gradient Directionality, PGD; theta mean 0.72, beta mean 0.62; Fig. 4E). For comparison, we conducted the same analysis in a lower-density recording (Fig. 4F-J). In this participant, beta waves dominated (Fig. 4H) with steeper phase gradient slopes indicating slower propagation speeds (theta mean 0.23 m/s, beta mean 0.96 m/s; Fig. 4I). Overall, travelling waves were again reliably detected (PGD theta mean 0.72, beta mean 0.71; Fig. 4J) and obeyed the same regularities as in the higher-density recording.

**Fig. 4.**
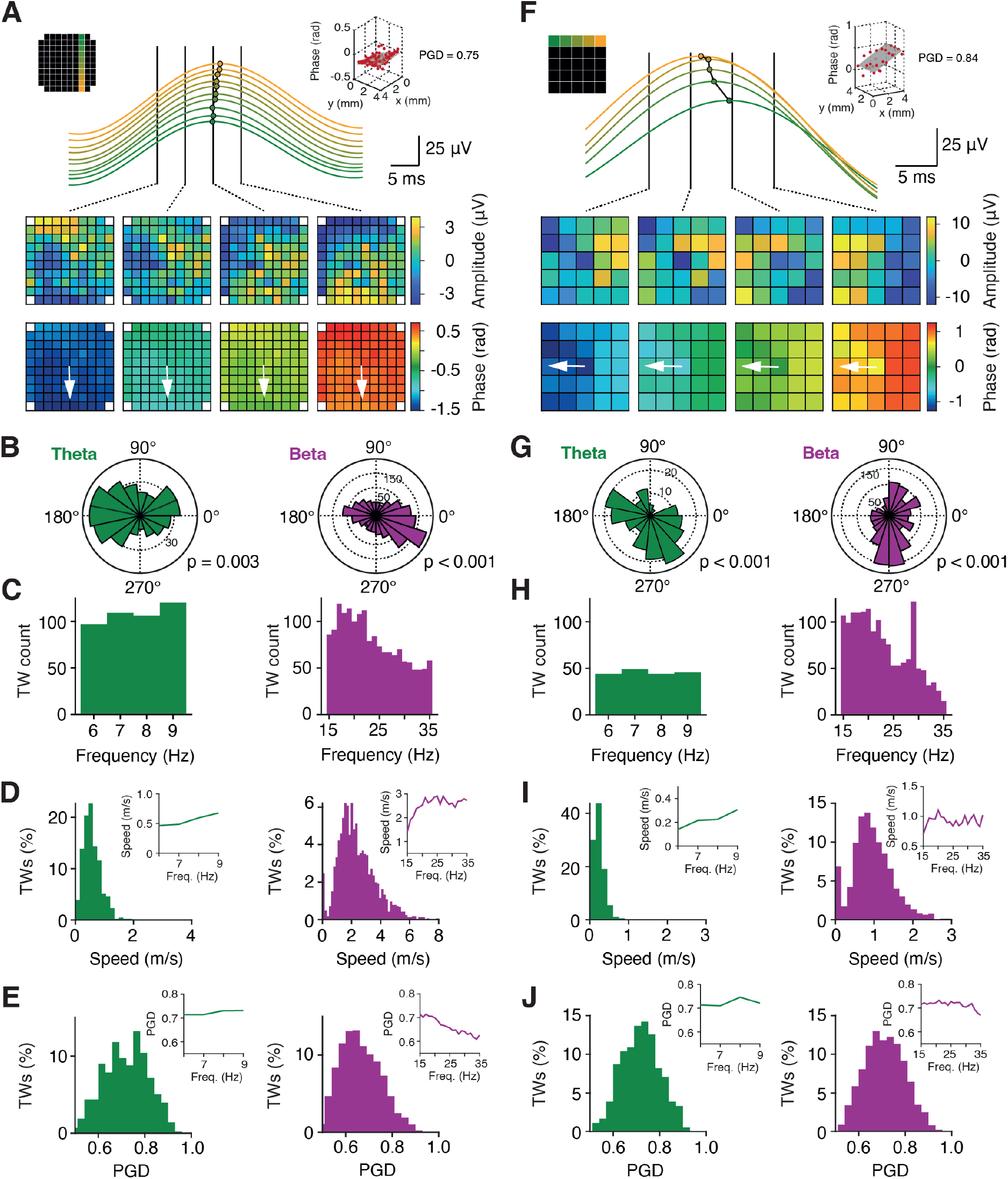
Propagation of waves of oscillatory activity across microelectrode arrays. (**A**) Example travelling wave recorded on a higher-density array. Top: peaks of LFP beta activity (20 ± 1.5 Hz) are temporally shifted across neighboring electrodes, illustrating the propagation of neural activity. Middle: demeaned LFP activity (amplitude) across the array at four example timepoints. Bottom: phase gradient across the array per timepoint. The arrow indicates the direction of wave propagation (from top to bottom). Inset: linear plane fitted to the phase gradient across the array at one example timepoint. (**B-E**) Distribution of travelling wave directions (B), count per frequency bin (C), speed (D) and plane model goodness-of-fit (PGD, E) in the theta (6 - 9 Hz, left) and beta (15 - 35 Hz, right) band in 500 ms epochs following the presentation of visual stimuli (sample numbers, see Fig. 5). Insets in (D) and (E) show frequency-resolved speed and PGD, respectively. p-values in (B) are given for Hodges-Ajne test for nonuniformity. (**F-J**) Same layout for travelling waves recorded on a lower-density array. PGD, phase gradient directionality.

**Fig. 5.**
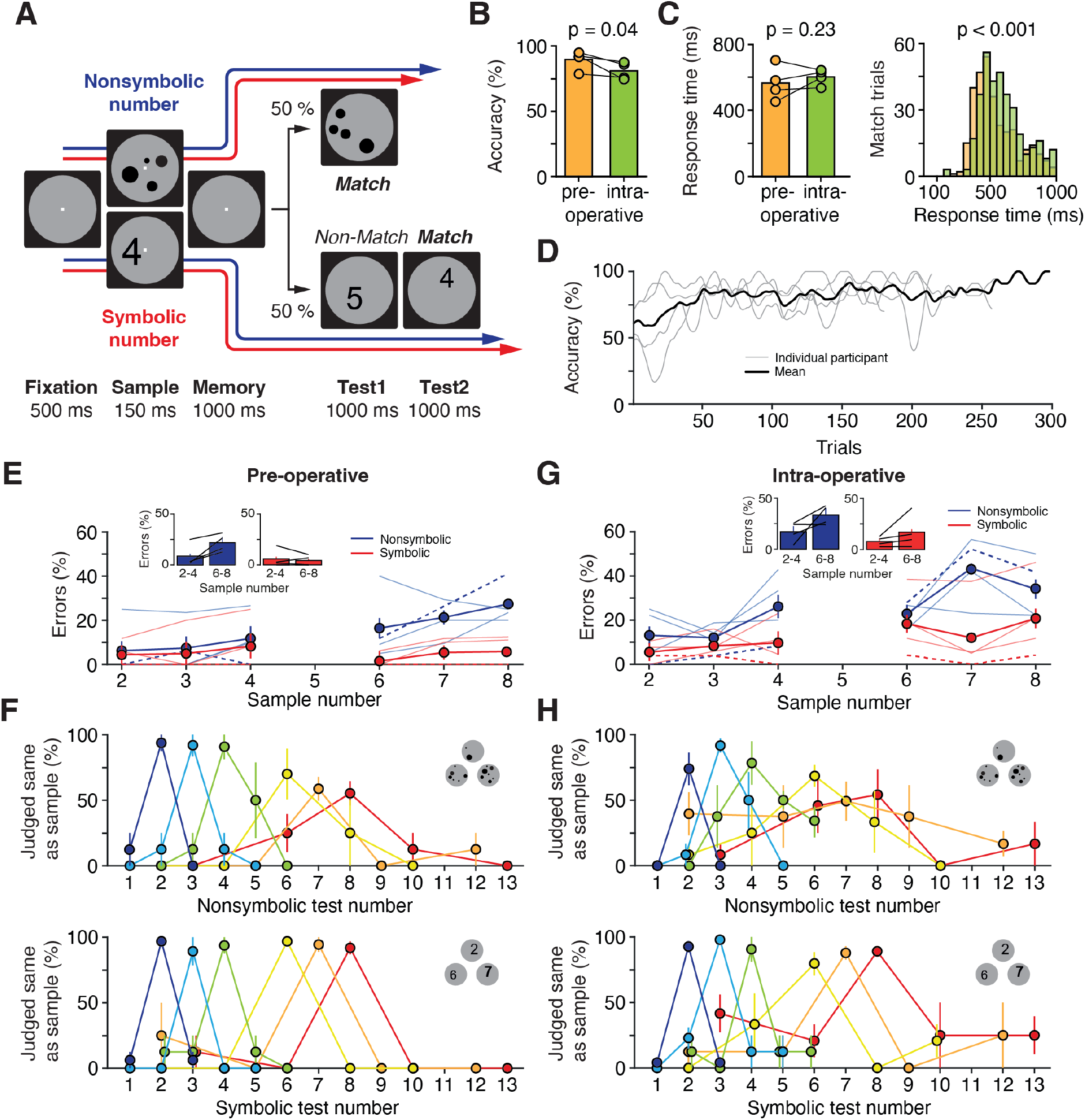
Preoperative and intraoperative cognitive performance in patients undergoing awake brain surgery. (**A**) Delayed-match-to-number task. Participants memorized the number of the sample stimulus and compared it to a subsequently presented test number. Trials were presented either in nonsymbolic notation (sets of dots, numerosities) or in symbolic notation (Arabic numerals). (B) Preoperative and intraoperative task performance (n = 4 participants; one-tailed t-test). (C) Preoperative and intraoperative response times in match trials on a per-participant basis (left) and pooled across trials (right) (one-tailed t-tests). (**D**) Time courses of intraoperative task performance across sessions. (**E**) Percentage of errors during preoperative behavioral testing plotted as a function of sample number and stimulus notation. Inset: performance pooled across small numbers (2-4) and large numbers (6-8). Error bars indicate SEM across participants. Dashed lines mark single-subject data for P10 (see Figs. 6, 7) (**F**) Preoperative behavioral tuning functions for trials with numbers presented in nonsymbolic and symbolic notation (top and bottom, respectively). Performance is shown for all sample-test-combinations. The peak of each curve represents the percentage of correct match trials, and other data points mark the percentage of errors in non-match trials. Error bars indicate SEM across participants. (**G**) Same layout as in (E) for intraoperative testing. (**H**) Same layout as in (F) for intraoperative testing.

In sum, our neurophysiological signal analysis showed that acquisition of multi-channel extracellular neuronal activity via intracortically implanted MEAs is feasible in the setting of awake brain surgery with its tight clinical and procedural constraints. Mesoscale network (LFP) activity for studying both local and propagating neuronal oscillations was obtained in high quality in every recording, while the extent of microscale spiking activity and yield of single units depended on the array configuration and favored the use of MEAs with increased electrode spacing.

### Probing higher cognitive functions in awake brain surgery

In parallel to neuronal data acquisition, we administered a task to the participants to probe the human number sense, a higher-level cognitive function of the parietal and (lateral) prefrontal association cortex that enables us to represent and manipulate abstract numerical categories (Nieder, 2016). The frontoparietal cortex has undergone disproportionate expansion in human evolutionary history, but is hardly ever targeted in single unit studies with DBS or epilepsy patients.

All six patients with recordings from either higher-density or lower-density arrays (Figs. 2 and 3) performed a delayed-match-to-sample task requiring them to memorize a visually presented sample number and compare it to a subsequently presented test number (Fig. 5A). Stimuli were presented either in nonsymbolic notation (sets of dots, numerosities) or in symbolic notation (Arabic numerals), allowing us to investigate the neuronal coding of and mapping between ‘non-verbal’ number, which animals have access to, and ‘verbal’ number, which is unique to humans. Four patients performed well in all conditions, whereas two patients (P07 and P09, higher-density arrays) did not exceed chance level in the nonsymbolic (dot) trials and were excluded from further analysis. There was only a small reduction in intra-operative response accuracy compared with pre-operative training levels (p = 0.04, one-tailed *t*-test; Fig. 5B) and a small increase in intra-operative response times (p = 0.23, one-tailed *t*-test per participant; p < 0.001, one-tailed Wilcoxon test with pooled trials; Fig. 5C). Following a brief ‘warm-up’ period, all patients maintained high performance levels throughout the recording session and completed between 200 and 300 trials (Fig. 5D).

The patients’ task performance was qualitatively very similar during pre-operative training and intra-operative recording and not distorted (compare Fig. 5E, F with Fig. 5G, H). Errors were more frequent during surgery, in nonsymbolic trials and for larger numbers (p_setting_ = 0.02, p_notation_ = 0.003, p_number_ = 0.01, 3-factorial ANOVA; Fig. 5E, G). Behavioral tuning functions (Fig. 5F, H) showed that participants correctly matched sample and test stimuli in particular for small numbers (peak of each curve), while accuracy dropped with increasing number. In non-match trials, the percentage of errors depended on the numerical distance between sample and test (distance effect; fewer errors for larger distances) and on the absolute magnitudes of the compared numbers (size effect; fewer errors for small numbers). Together, these results show that all key behavioral signatures of numerical cognition were captured by the task administered to the participants.

### Human neuronal coding of number at the micro- and mesoscale level

Extracellular recordings in the non-human primate frontoparietal cortex suggest that single units tuned to individual numerosities give rise to numerical cognitive abilities (Jacob et al., 2018; Jacob & Nieder, 2014; Nieder et al., 2006). The human neuronal code for number in these brain areas, however, is not known. Leveraging the flexibility in array placement and high-quality data obtained with MEA recordings from open craniotomies, we illustrate here a potential application of this method by exploring - in parietal cortex (inferior parietal lobule, IPL) of an example participant (P10) - the neuronal correlates of the human number sense at the single-neuron and neuronal network level.

In nonsymbolic trials, an example single unit strongly increased its firing rate after presentation of the sample stimulus (Fig. 6A, left). The increase was graded and a function of sample numerosity with peak activity for 7 and 8 dots. This unit’s firing rates were smaller and more transient in trials with symbolic number, but showed a similar graded response (Fig. 6A, right). Average firing rates in the 500 ms epoch following sample presentation confirmed significant tuning to nonsymbolic number, but failed to reach significance in symbolic trials due to the distinct temporal activity profile (Fig. 6B). Thus, this single unit carried information (ω^2^ percent explained variance) about sample notation and numerosity (Fig. 6C). Similar responses were found in a different example single unit recorded on a neighboring electrode (Fig. 6D-F). An example multi-unit measured on a different electrode of the same array was tuned to nonsymbolic number 1 (Fig. 6G, left). This unit also showed a congruent response in trials with symbolic numbers, albeit with distinct dynamics and a more categorical coding of small versus large numbers (Fig. 6G, right and Fig. 6H, I).

**Fig. 6.**
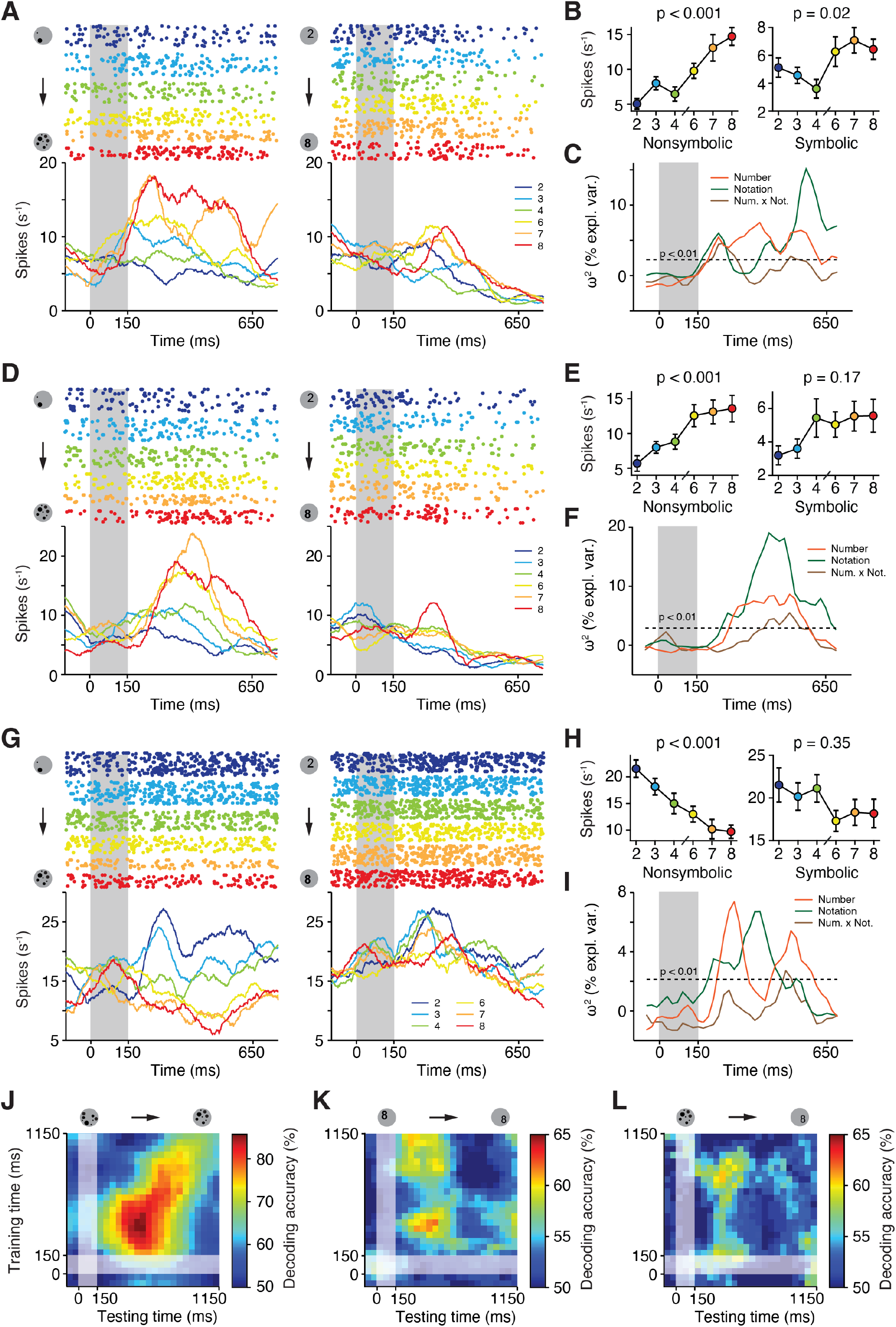
Single unit and neuronal population coding of nonsymbolic and symbolic number. (**A**) Spike raster plots and spike-density histograms (smoothed using a 150 ms Gaussian window) for an example single unit recorded in the inferior parietal lobe. Trials are sorted by sample numerosity and by stimulus notation (left: nonsymbolic, right: symbolic). Sample presentation is highlighted. (**B**) Firing rate of the neuron in (A) in the 500 ms epoch following presentation of nonsymbolic and symbolic sample numerosities (left and right, respectively; one-factorial ANOVA). (**C**) Sliding-window ω^2^ percent explained variance (two-factorial ANOVA) quantifying the information about sample number and notation as well as their interaction contained in the firing rate of the neuron in (A) in correct trials. Dashed line marks the significance threshold (p = 0.01; shuffle distribution). (**D-F**) Same layout as in (A-C) for a different single unit recorded on a neighboring channel on the same microelectrode array. (**G-I**) Same layout as in (A-C) for a multi-unit recorded on a neighboring channel on the same microelectrode array. (**J**) Cross-temporal LDA decoding of nonsymbolic number (small, i.e. 2-4, versus large, i.e. 6-8) in the 1000 ms memory epoch following sample presentation using spiking activity (multi-units) on all channels of the microelectrode array. Sample presentation is highlighted. (**K**) Same layout as in (J) for symbolic number. (**L**) Same layout as in (J) for cross-notation decoding. The decoder was trained in trials with nonsymbolic numerosities and tested in trials with symbolic numerosities.

To provide a population-wide perspective on number coding, we trained a linear discriminant analysis (LDA) decoder to separate small from large numerosities using the entire spiking activity recorded across the array (Fig. 6J-L). In trials with nonsymbolic number, decoding accuracy was high and peaked (86 %) after sample presentation, matching the single unit responses. Cross-temporal training and decoding showed a dynamically evolving code across the memory delay with reduced off-diagonal accuracy (Fig. 6J). In trials with symbolic number, decoding was less accurate (62 % peak) and only possible in the first half of the memory delay, again matching single unit responses (Fig. 6K). The results of cross-notation decoding (training on nonsymbolic number, testing on symbolic number) were qualitatively similar with decoding accuracy bounded by the weaker coding of symbolic number compared to nonsymbolic number (Fig. 6L).

We then directly compared the microscale neuronal activity elicited during the task with mesoscale network responses. At the same electrode on which the number-tuned single unit shown in Fig. 6A-C was recorded, LFP power varied strongly with sample number and notation (and their interaction) in particular in the gamma band (45 - 100 Hz; ω^2^ percent explained variance; Fig. 7A). However, in contrast to the early changes in spiking activity, sample selectivity measured by LFPs increased only 150 ms after sample offset (compare e.g. Fig. 7A left with Fig. 6A left). In the 500 ms epoch following sample number presentation, gamma power increased monotonically with numerosity in nonsymbolic trials, but did not vary with symbolic number (p < 0.001 and p = 0.46, respectively, one-factorial ANOVA; Fig. 7B top). On two neighboring channels (same electrodes on which units shown in Fig. 6D-F and Fig. 6G-I were recorded) a qualitatively similar pattern was found (p < 0.001 and p = 0.02, respectively, one-factorial ANOVA; Fig. 7C, D top), albeit with a clear spatial gradient. Beta responses, in contrast, were spatially more uniform, underscoring the local nature of gamma activity and the potentially distinct functional reach of the analyzed frequency bands (Fig. 7B-D bottom). Of note, while not all units in Fig. 6 were tuned to the same preferred number, LFP power scaled uniformly with numerosity across electrodes (compare Fig. 6G left with Fig. 7D top). Analysis of propagating oscillatory activity across the array also showed that, at equal strength, travelling waves were faster for larger numerosities (Fig. 7E).

**Fig. 7.**
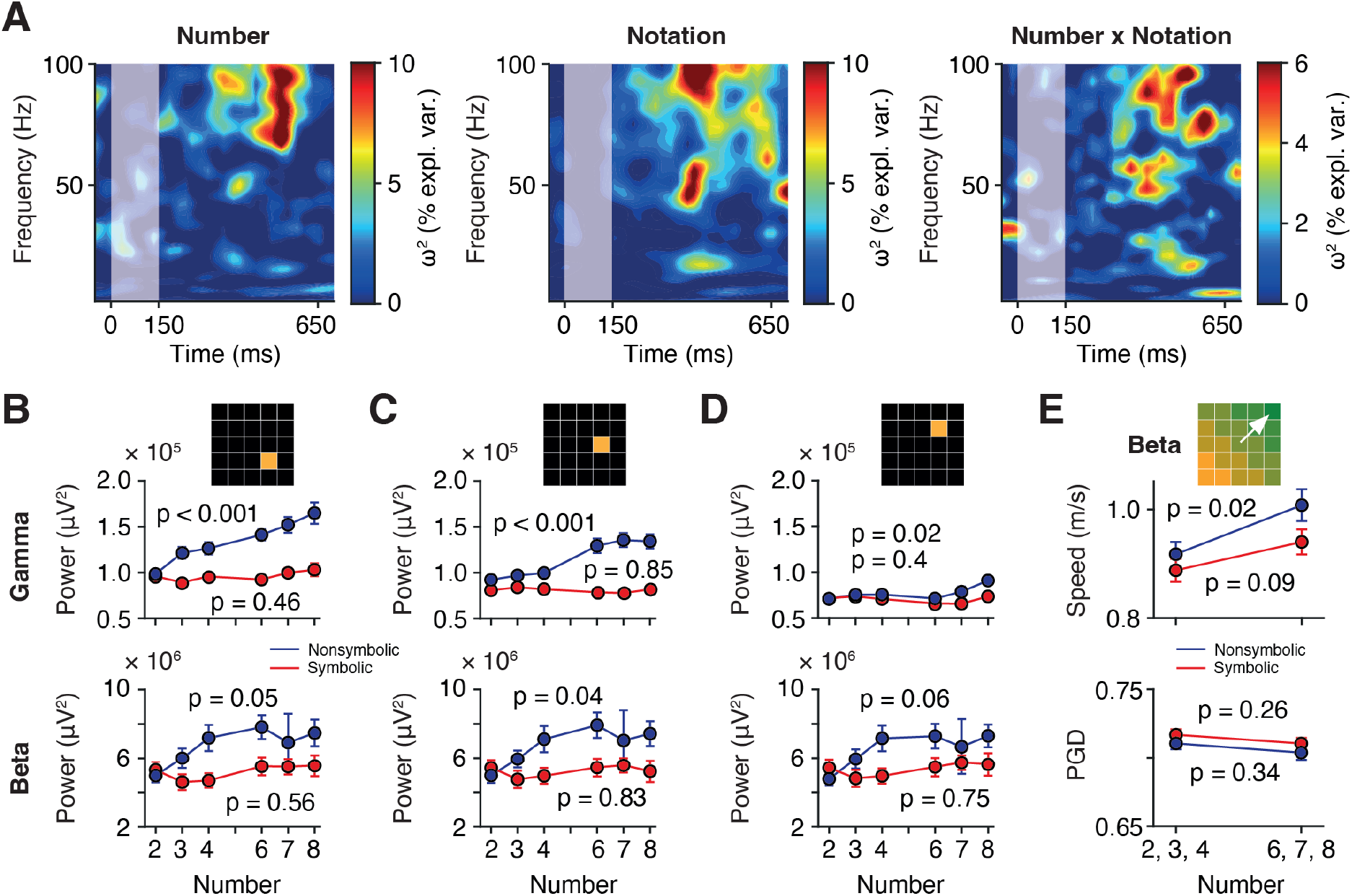
Local and propagating oscillatory neuronal activity during number coding. (**A**) Sliding-window ω^2^ percent explained variance (two-factorial ANOVA) quantifying the information about sample number (left) and notation (middle) as well as their interaction (right) contained in the LFP power spectrum of an example single channel on a lower-density array (same channel as in Fig. 6A-C) in correct trials. Sample presentation is highlighted. (**B**) LFP power in the gamma (45 - 100 Hz, top) and beta (15 - 35 Hz, bottom) band in the 500 ms epoch following sample number presentation as a function of sample number in nonsymbolic and symbolic notation. Same channel as in (A). p-values are given for one-factorial ANOVA. (**C**) Same layout as in (B) for a neighboring single channel. (D) Same layout as in (C) for a neighboring single channel. (**E**) Speed (top) and goodness-of-fit (PGD, bottom) of LFP beta band travelling waves propagating across the array in the 500 ms epoch following sample number presentation for small (2-4) and large (6-8) numbers in nonsymbolic and symbolic notation. p-values are given for one-factorial ANOVA.

Our proof-of-concept results suggest that, first, the human parietal cortex harbors single units that are tuned to number, establishing a previously missing link to the non-human primate animal model. Second, at the single-neuron level, nonsymbolic set sizes are coded with graded and continuous responses, displaying no sign of neuronal subitizing. A well-studied behavioral signature of the approximate number system, subitizing denotes the accurate apprehension of small numbers of items at a glance (evidenced by a disproportionate increase in errors for larger numerosities in nonsymbolic, but not symbolic notation; single-subject data for P10 [dashed lines] in Fig. 5E, G) and is thought to indicate different representational systems for small and large quantities (Piazza et al., 2011). Our findings are not compatible with this hypothesis and rather argue that the representation of small and large quantities emerges from a single system (Cheyette & Piantadosi, 2020). Third, symbolic numbers are coded with distinct temporal dynamics and more categorical responses than nonsymbolic quantities, in line with recent findings in the human MTL (Kutter et al., 2018). However, the number code partially generalizes across notations with number-congruent responses for nonsymbolic and symbolic stimuli. Fourth, spiking activity and oscillatory activity reflect distinct aspects of numerical information processing in the local microcircuit, with LFPs possibly capturing in particular the network’s load-dependent activity state.

## Discussion

We found that intracortically implanted MEAs are suitable for acute recordings of human brain activity at both meso- and microscale resolution (Figs. 2–4). All arrays acquired LFPs (synaptic network activity) with high fidelity. Increasing the interelectrode spacing also allowed us to record responses from populations of single units. The devices can be used in awake surgeries with large open craniotomies, providing broad access to the cortex (Fig. 1) in patients who achieve close to normal levels of cognitive performance (Fig. 5). We illustrated a potential application by exploring the neuronal correlates of human numerical cognition in parietal cortex (Figs. 6, 7), a brain region that is typically inaccessible in DBS or epilepsy surgery, i.e. in procedures that so far have produced the vast majority of intracranial data tapping into the neuronal underpinnings of human cognitive functions.

We believe the comparative ease with which MEA recordings can be introduced into the operating room and incorporated into established neurosurgical procedures to be their greatest advantage. Positioning of the array and implantation can be completed within ten minutes. After insertion, the arrays ‘float’ on cortex. No extra manipulators or electrode holders are required (Chung et al., 2022; Paulk et al., 2022). The arrays readily follow brain movements, yielding stable recordings without the need for additional mechanical stabilization (Jamali et al., 2019; Jamali et al., 2021). Slight shifts of the skull in awake participants and above all vertical displacements of the cortex during brain pulsations pose a major challenge when externally secured probes are used that occupy a different spatial reference frame than the tissue they record from, necessitating elaborate post-acquisition motion correction (Chung et al., 2022; Paulk et al., 2022). Furthermore, penetrating MEAs are robust, have a well-documented safety profile and are used with equipment that has been validated for sterilization and re-use. There is no risk of shank breakage, no inadvertent deposition of electrode material in brain tissue, and no need to perform piotomies to allow entry of the device into cortex as with more delicate (e.g. Neuropixels) probes (Chung et al., 2022; Paulk et al., 2022). Good grounding could be reliably achieved either by anchoring the pedestal to the skull or by establishing a strong connection to the head frame. Both configurations were effective in our experience and sufficient to reduce electrical hum and noise to levels that enable high-quality extracellular recordings despite an environment full of potential sources of interference. We did not find it necessary to turn off suction, lighting, warming blankets or any other piece of medical equipment during recording.

The arrays’ grid-like electrode arrangement allows for dense sampling of neuronal activity in the horizontal plane, i.e. from a patch of cortex. There is rapidly mounting interest in the mechanisms by which propagating neuronal activity, e.g. in form or travelling waves (Fig. 4), mediates intercortical information transfer (Bhattacharya et al., 2022; Das et al., 2022; Rubino et al., 2006; Sato et al., 2012; Takahashi et al., 2011; Zhang & Jacobs, 2015; Zhang et al., 2018). In contrast to microwire bundles with their irregularly placed electrode tips or linear probes that record from one single cortical column, MEAs with their well-defined planar geometry are ideally suited to address such questions. Spatial coverage may be extended even further by the addition of ECoG grids, which can be placed directly on top of MEAs, or intracranial stereo EEG leads (Chiang et al., 2020; Tong et al., 2021; Vaz et al., 2020). Lastly, using MEAs in open craniotomy surgeries where the implanted tissue is resected (as in our participants) opens up the possibility of complementing the *in vivo* recordings with *in vitro* physiological or histological analyses to explore structural-functional relationships in neural circuit organization (Loomba et al., 2022).

MEAs with increased interelectrode spacing (25 channels) recorded on average more than one well-isolated single unit per channel (Fig. 3). Per patient and recording session, this yield is similar to semi-chronic recordings in epilepsy patients (2 to 3 neurons per microwire bundle with up to 10 bundles implanted per patient (Fu et al., 2022; Kutter et al., 2018)). Acute DBS recordings from prefrontal cortex (10 to 20 neurons per participant (Jamali et al., 2019; Jamali et al., 2021)) or midbrain structures (fewer than 10 neurons per participant (Zaghloul et al., 2009; Zaghloul et al., 2012)) yield less. Efforts are currently underway to establish acute intracranial recordings with high-density linear probes (Neuropixels), which have been reported to pick up between several tens of neurons in open craniotomies (Chung et al., 2022) to a few hundred units in DBS burr holes (Paulk et al., 2022). Critical technical challenges are still to be met, but these probes could eventually provide a valuable addition to the armamentarium of intraoperative recording devices from which the neurophysiologist and neurosurgeon can chose depending on the particular research question and clinical setting.

The arrays’ geometrical configuration was a crucial determinant of spiking activity SNR (Fig. 2). This is likely a consequence of the electrodes’ comparatively large footprint (thickness 180 - 200 μm near the base), the main disadvantage of the MEAs used in this study. Lower-density arrays produce less cortical trauma, thereby increasing the chances of measuring single unit activity shortly after array insertion. Our histological analyses showed microhemorrhages in some (Fernandez et al., 2014; House et al., 2006), but not all implantations of standard 96 channel arrays. Cortical neuronal ‘stunning’ might therefore be an important reason for the very low single unit yield in higher-density arrays. Fittingly, unit activity in our recordings only appeared after several minutes and continued to develop until data acquisition began when the patient was fully awake, a time period significantly longer than recently reported for thinner linear probes (Chung et al., 2022; Paulk et al., 2022). A second limitation of the described setup is the difficulty in precisely controlling pneumatic array insertion. Whether the inserter wand is stabilized by a dedicated holder or manually (we preferred the latter to expedite implantation), the inherent variability in inserter positioning will significantly affect the forces that the electrode pad experiences during implantation, much unlike micromanipulator-controlled implantations of e.g. linear probes. Imperfect alignment of the inserter with the array could disproportionately impact implantations of higher-density arrays and in older patients (Fernandez et al., 2014), where optimal forces are required to overcome the increased resistance to insertion from the pial meninges and brain tissue. We found it best to place the inserter into direct contact with the array, applying very gentle downward pressure to eliminate dead space between the electrode tips and cortical surface (Fig. 1). This approach resulted in complete array insertions and reproduceable signals for both higher-density and lower-density arrays (Fig. 2).

High-volume recordings are necessary to accelerate progress in our understanding of the neuronal basis of human brain functions. Awake surgeries for tumor resection are performed at many medical centers. We have shown here that these procedures are as suitable for acquiring cellular resolution data from the human brain as DBS or epilepsy surgeries. As any other probe in the expanding palette of multichannel recording devices (Chung et al., 2022; Paulk et al., 2022), intracortical MEAs do not promise a fail-safe or turn-key solution. However, the technology is more mature and more lenient in the intraoperative setting where clinical constraints considerably limit options for optimizing the recording setup and neuronal signal quality. Once mastered, it can also be effectively put to use in chronic (e.g. BCI) applications where MEAs represent the gold-standard for intracranial sensors. Human single-unit recordings are multidisciplinary endeavors, for which all stakeholders must advance beyond their comfort zones. The methods we describe here can stimulate productive collaborations between neuroscientists and clinicians and propel forward the exploration of the unique neural computations performed by the human brain.

## Materials and Methods

### Experimental design

We included 13 participants in this study with intracerebral tumors (mainly glioblastoma) referred to our department for surgical resection (Table 1). All study procedures were conducted in accordance with the Declaration of Helsinki guidelines and approved by institutional review board (IRB) of the Technical University of Munich (TUM) School of Medicine (528/15 S). Participants were enrolled after giving informed consent. The scientific aims of this study had no influence on the decision to operate. With the exception of array implantation, the course of the surgery was not altered.

### Multielectrode arrays and implantation procedure

Per participant, one Neuroport IrOx planar multielectrode array (Blackrock Neurotech) was implanted. In nine patients, we implanted the standard array with 96 wired (active) electrodes on a 10×10 grid (1.5 mm electrode length, interelectrode spacing 400 μm). In four patients, we implanted a custom array with 25 channels, which was produced by removal of every second row and column from the standard array (interelectrode spacing 800 μm; Fig. 2c). The array’s pedestal was first anchored to the skull adjacent to the craniotomy. The array was then positioned on the cortical surface of the to-be-implanted gyrus guided by MRI-neuronavigation (Brainlab, Germany). Care was taken to avoid prominent vascular structures, which in some cases prompted us to deviate from the preoperatively determined implantation site by a few millimeters. References wires were inserted under the dura.

The array was implanted pneumatically following the manufacturer’s guidelines (Blackrock Neurotech). We found that introducing a dedicated external wand holder was inconvenient, and that positioning of the holder unnecessarily prolonged the implantation procedure. We therefore secured the wand manually such that it touched the array’s dorsal pad and brought the electrode tips into contact with the pia. Insertion was performed with a single pulse (20 psi, pulse width 3.5 ms). We did not systematically explore different insertion pressure or pulse width settings. The array was then covered with saline irrigated strips and left to settle as the patient was allowed to awake from anesthesia.

All equipment in contact with the patient (inserter wand, trigger, tubing, headstages, cabling) was re-sterilized (Steris V-Pro) and used in multiple surgeries.

In all participants, the implantation site was chosen to lie within the resection area surrounding the tumor. In some cases, however, intraoperative evaluation determined that the implanted tissue could not be safely resected, so that the array was removed from the brain tissue prior to closure of the dura and the craniotomy. In three participants (P01, P02 and P03), the resected implantation region was formalin-fixed with the array *in situ* and processed further for histological analysis (hematoxylin eosin staining).

Cortical surfaces were reconstructed from individual participants’ structural MRI using BrainSuite (Shattuck & Leahy, 2002). The implantation site was marked manually, guided by intraoperative neuronavigation data and photographic documentation. Individual MRI scans were then normalized to the MNI-152 template in SPM12 (Wellcome Center Human Neuroimaging). The macroanatomical cortical area corresponding to the implantation site was determined using the JuBrain SPM anatomy toolbox (Forschungszentrum Jülich).

### Neurophysiological recordings

We recorded intraoperative neuronal data in eight participants. Extracellular voltage signals were acquired using either analog patient cable headstages in combination with a front-end amplifier (P04, P05, P06, P07 and P09) or digital Cereplex E128 headstages connected to digital hubs (P10, P11 and P13) as part of a 128-channel NSP system (NeuroPort Biopotential Signal Processing System, Blackrock Neurotech). Settings for signal amplification, filtering and digitization were identical in both setups (high-pass 0.3 Hz, low-pass 7.5 kHz, sampling rate 30 kHz, 16-bit resolution).

We did not find it necessary to switch between the two reference wires, both of which provided high-quality reference signals in all cases. However, particular attention was paid to achieving a strong ground connection via the pedestal. Long skull screws (6 mm) in combination with intermittent irrigation of the pedestal’s base where it contacted the skull produced the best results. Impedances were checked after array implantation and in most surgeries were initially higher than the upper bound of the normal range (80 kΩ for IrOx electrodes), but continued to normalize over the course of several tens of minutes. We attributed this to improving electrical conductivity at the pedestal-skull interface. Additional ground connections were not necessary and could even contaminate signals if placed badly (e.g. subdermal needles in the vicinity of musculature).

### Behavioral task and stimuli

Six participants performed a delayed-match-to-number task during neuronal recording. MonkeyLogic 2 (NIMH) running on a dedicated PC was used for experimental control and behavioral data acquisition. Behavioral time stamps were transmitted to the NSP system for parallel logging of neuronal data and behavioral events.

We familiarized participants with the task ahead of the surgery and allowed them to complete multiple training trials. Participants viewed a 12” monitor positioned 40 - 50 cm in front of them. They were instructed to maintain eye fixation on a central white dot and pressed a button on a hand-held device to initiate a trial. Stimuli were presented on a centrally placed gray circular background subtending approx. 9,4 ° of visual angle. Following a 500 ms pre-sample period, a 150 ms sample stimulus was shown. In nonsymbolic trials, 2, 3, 4, 6, 7 or 8 randomly arranged black dots specified the corresponding numerosity. In symbolic trials, black Arabic numerals (Arial, 40 - 56 pt) were shown. The participants were required to memorize the sample number for 1,000 ms and compare it to the number of dots (in nonsymbolic trials) or the Arabic numeral (in symbolic trials) presented in a 1,000 ms test stimulus. If the quantities matched (50 % of trials), participants released the button (correct Match trial). If the quantities were different (50 % of trials), the participants continued to push the button until the matching quantity was presented in the subsequent image (correct Non-match trial). Match and non-match trials and nonsymbolic and symbolic trials were pseudo-randomly intermixed.

New stimuli were generated for each participant and recording. Low-level, non-numerical visual features could not systematically influence task performance (Jacob et al., 2018): in half of the nonsymbolic trials, dot diameters were selected at random. In the other half, dot density and total occupied area were equated across stimuli.

### Behavioral performance

Behavioral tuning functions were used to describe the percentage of trials (y axis) for which a test stimulus (x axis, units of numerical distance to sample number) was judged as being equal in number to the sample. A numerical distance of 0 denotes match trials; the data point represents the percentage of correct trials. As the numerical distance increases, there is less confusion of the test with the sample number; the data points represent the percentage of error trials. Tuning curves were calculated separately for trials with nonsymbolic stimuli and for trials with symbolic stimuli.

### Spiking activity and single unit quality metrics

Raw signals were filtered (250 Hz high-pass, 4-pole Butterworth), and spike waveforms were manually separated from noise using Offline Sorter (Plexon). Signal-to-noise ratio (SNR) was calculated as

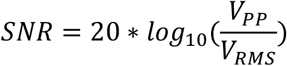

where V_pp_ is the mean peak-to-peak spike amplitude of a given channel and V_RMS_ is the root-mean-square (RMS) voltage

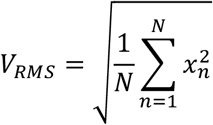

with x_n_ being individual voltage values (Fig. 2D top). Spike SNR was calculated across the entire recording session (Fig. 2D bottom) or in sliding windows (Fig. 2E; 60 s bins, 30 s steps).

Thresholded waveforms were manually sorted into clusters of single units (Offline Sorter). We estimated the rate of false negatives (missed spikes) by fitting a gaussian to the distribution of spike troughs (Fig. 3D). Autocorrelograms (Fig. 3E) were calculated by shifting a unit’s spike train in steps of 1 ms over a range of 1 to 25 ms. To determine the percentage of outlier spikes (Fig. 3G) (Meirhaeghe et al., 2021), each spike was considered as a point on a 2D plane spanned by the first two principal components that were used for spike sorting. For each spike, the Mahalanobis distance to the corresponding cluster’s average waveform was calculated. A chi-square distribution was then fitted to the distribution of distances (Hill et al., 2011). If the likelihood of a given spike to belong to this distribution was lower than a fixed threshold (the inverse of the total number of spikes in the given cluster), it was considered an outlier spike.

### Local field potentials and quality metrics

Data was processed using the FieldTrip toolbox (Oostenveld et al., 2011). Raw signals were filtered (1.5 Hz high-pass, 1-pole Butterworth; 250 Hz low-pass, 3-pole Butterworth), and line noise was removed (2-pole Butterworth band-stop filters of ± 0.2 Hz at 50 Hz and harmonics). LFP traces were then visually inspected for large-amplitude artefacts, which were excluded from further analysis.

Spectral transformation was performed with the additive superlet method (Moca et al., 2021). SNR was calculated in sliding windows (60 s bins, 30 s steps) and then averaged across windows for the session-SNR (Fig. 2H bottom) or presented as time-resolved data (Fig. 2I). For each bin and channel, states of high and low LFP activity were identified and used for signal and noise estimators, respectively (Fig. 2H top) (Compte et al., 2008; Suarez-Perez et al., 2018). High and low activity states were derived from the smoothed LFP amplitude envelope (100 ms averaging window) obtained through complex Hilbert transform. Any timepoints of the smoothed envelope that fell outside of three standard deviations of its distribution were marked as artefacts and automatically assigned to the noise intervals. The mean of the smoothed envelope, excluding artefact timepoints, served as a detection threshold for high activity states. Thus, epochs of the smoothed envelope surpassing the threshold for at least 400 ms were considered states of high activity, whereas all others counted as low activity states (Compte et al., 2008). SNR was then calculated as

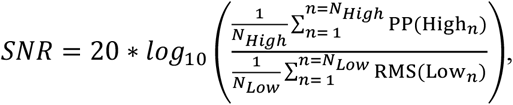

where N_High_ and N_Low_ are the number of high or low activity states, respectively, PP (peak-to-peak amplitude) is the difference between the highest and lowest voltage reading during a given high activity state and RMS is

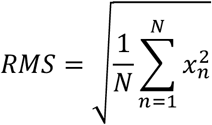

with x_n_ being individual voltage values of an interval of low activity.

The Power-Spectral-Density (PSD) was calculated using Welch’s method. Specifically, across five minutes of the recording (0:30 to 5:30 min), modified periodograms in 3-s bins (smoothed using a Hamming window) with 50 % overlap were obtained by Fast Fourier transform (FFT) and averaged (Zilio et al., 2021).

### Travelling waves

We assumed the simplest form of travelling waves, a planar wave with linear phase gradient (Rubino et al., 2006). First, zero-phase bandpass filters (± 1.5 Hz) were applied for each frequency of interest (theta: 6 to 9 Hz; beta: 15 to 35 Hz, in steps of 1 Hz) and every channel. We then applied the Hilbert transform (Hlb) to the resulting signal (V) to obtain the instantaneous phase φ(x,y,t) of each time point (t) and channel position (x,y)

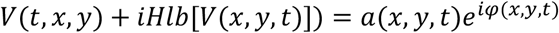

Instantaneous phases were unwrapped and de-noised (Woods, 2011). Next, a plane model was fit to the data using linear regression. The plane was modelled as

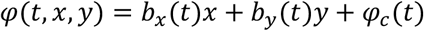

With b_x_(t) and b_y_(t) being the slope of the plane in the x-direction and y-direction at time t, respectively, and φ_c_(t) the constant phase shift at time t. The model’s goodness-of-fit was expressed by the Phase-Gradient Directionality (PGD) (Rubino et al., 2006). PGD is the Pearson correlation between the predicted and actual phase and is given by

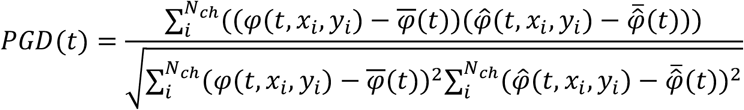

with 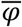 being the average and 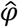 the predicted phase.

When zero fell outside the 99^th^ percentile of at least one of the coefficients’ b_x_ or b_y_ confidence intervals and PGD was bigger than 0.5, a moment in time was considered for travelling wave-like activity (Rubino et al., 2006). The direction (Woods, 2011) and speed (Rubino et al., 2006) of the travelling wave-like activity were then calculated as

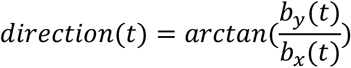

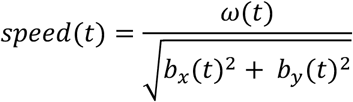

with ω(t) being the instantaneous angular velocity.

A travelling wave epoch was defined by non-zero slopes in the phase gradient with a PGD > 0.5 for a minimum length of 5 ms and a maximal average change in direction of 3 deg/ms. Polar distributions (10° bins) that showed a second peak reaching 25 % or more of the distribution’s modal value and that significantly differed from uniformity (Hodges-Ajne test) were considered bidirectional.

### Neuronal information

To quantify the information about sample number and notation that was carried by a neuron’s spiking rate, we used the ω^2^ percent explained variance measure (Jacob & Nieder, 2014). ω^2^ reflects how much of the variance in a neuron**’**s firing rate can be explained by a given factor. It was calculated in sliding windows (100 ms bins, 20 ms steps) using

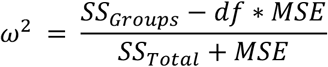

where the individual terms are derived from a two-way categorical ANOVA: *SS_Groups_* denotes the sum-of-squares between groups (numbers), *SS*_*Tota*l_ the total sum-of-squares, *df* the degrees of freedom, and *MSE* the mean squared error. The number of trials in each group was balanced. Balancing was accomplished by stratifying the number of trials in each group to a common value: A random subset of trials was drawn (equal to the minimum trial number across groups) and the statistic was calculated. This process was repeated 25 times, and the overall statistic was taken to be the mean of the stratified values. Significance thresholds were determined by randomly shuffling the association between spiking rates and trial type (number and notation) during the pre-sample epoch (500 ms). This process was repeated 1,000 times, and the significance threshold was set to the 99^th^ percentile of the cumulative distribution (p < 0.01).

For task information contained in LFPs, we calculated ω^2^ in sliding windows (5 ms bins, 0.25 ms steps, 1 Hz bins, 1 Hz steps) using spectral power derived as described above.

### Linear discriminant analysis

Unsorted (multi-unit) spikes were aggregated into firing rates using Gaussian windows with 50 ms sigma and 50 ms step size. Trials were grouped for small numbers (2, 3, 4) and large numbers (6, 7, 8). A procedure of 7-fold cross validation with 7 repetitions was used, resulting in 49 training and testing set pairs. At every time step, an LDA decoder (Scikit-learn package in Python) was trained on the activity of the current time step in the training set and tested on all the time steps in the testing set in order to investigate how well the code generalizes across different timesteps. Decoding accuracy is given as the average across test trials. LDA finds the component that maximizes the Mahalanobis distance between the centroids of small and large number classes. The algorithm assumes equal within-class covariance in different classes. Shrinkage of the empirical covariance matrix was applied by averaging the empirical covariance matrix with a diagonal matrix, discounting the spurious covariation between units. The amount of shrinkage was determined by the Ledoit-Wolf lemma (Ledoit & Wolf, 2004).

### Statistical analysis

All data analysis was performed with MATLAB (Mathworks) and Python.

## Acknowledgements

This study was supported by grants from the German Research Foundation (DFG JA 1999/5-1), the European Research Council (ERC StG MEMCIRCUIT, GA 758032) and the Else Kröner-Fresenius Foundation (TUM Doctorate Program Translational Medicine) to S.N.J., a grant from the Technical University of Munich (TUM Innovation Network Neurotech) to S.N.J. and J.G., and by a grant from the German Research Foundation to J.G. (GE 3008/3-1). We would like to thank Doris Droese for technical assistance with intraoperative recordings and Sandra Baur, Claire Delbridge and Friederike Liescher-Starnecker for preparing histological sections. We are especially indebted to our patients for their willingness to participate in this research.

---

**Fig. S1.**
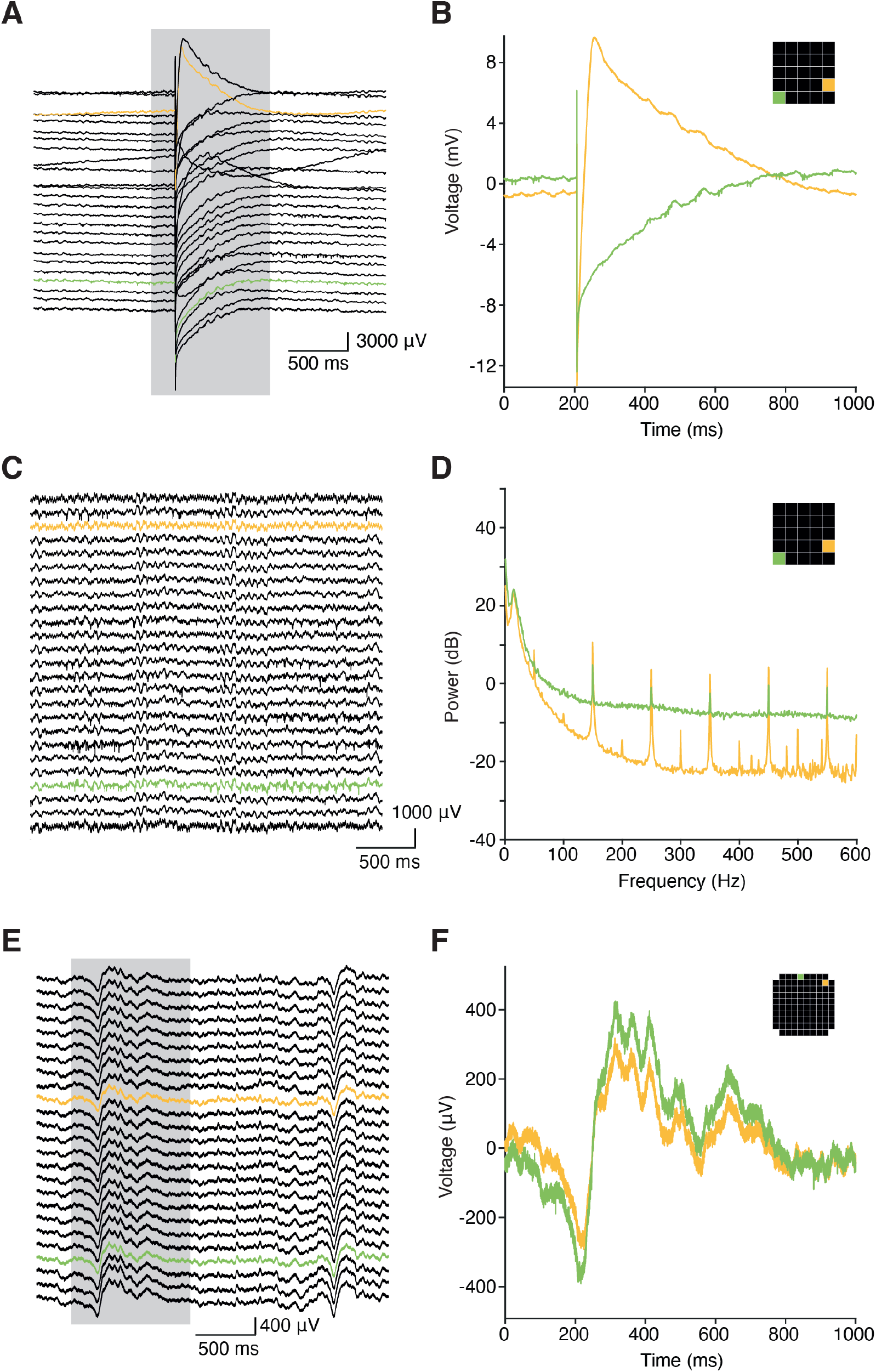
Example electrical artefacts during intraoperative recording. (**A, B**) Single large-amplitude electrode ‘pop’ with prolonged voltage settling time in a lower-density array recording. Note the voltage scale and compare to subsequent panels. Two representative channels are highlighted in (B) together with their location on the MEA grid (inset). (**C, D**) Line noise (50 Hz) and its harmonics in the same recording as in (A, B). (**E, F**) Contamination of the ground in a higher-density array recording by frontal facial and ocular muscle activity leading to intermittent slow artefacts.

## References

Aflalo, T., Kellis, S., Klaes, C., Lee, B., Shi, Y., Pejsa, K., Shanfield, K., Hayes-Jackson, S., Aisen, M., Heck, C., Liu, C., & Andersen, R. A. (2015). Decoding motor imagery from the posterior parietal cortex of a tetraplegic human. Science, 348(6237), 906–910. https://doi.org/10.1126/science.aaa5417

Bhattacharya, S., Brincat, S. L., Lundqvist, M., & Miller, E. K. (2022). Traveling waves in the prefrontal cortex during working memory. PLoS Comput Biol, 18(1), e1009827. https://doi.org/10.1371/journal.pcbi.1009827

Cash, S. S., & Hochberg, L. R. (2015). The emergence of single neurons in clinical neurology. Neuron, 86(1), 79–91. https://doi.org/10.1016/j.neuron.2015.03.058

Chen, X., Wang, F., Fernandez, E., & Roelfsema, P. R. (2020). Shape perception via a high-channel-count neuroprosthesis in monkey visual cortex. Science, 370(6521), 1191–1196. https://doi.org/10.1126/science.abd7435

Cheyette, S. J., & Piantadosi, S. T. (2020). A unified account of numerosity perception. Nat Hum Behav, 4(12), 1265–1272. https://doi.org/10.1038/s41562-020-00946-0

Chiang, C. H., Won, S. M., Orsborn, A. L., Yu, K. J., Trumpis, M., Bent, B., Wang, C., Xue, Y., Min, S., Woods, V., Yu, C., Kim, B. H., Kim, S. B., Huq, R., Li, J., Seo, K. J., Vitale, F., Richardson, A., Fang, H., … Viventi, J. (2020). Development of a neural interface for high-definition, long-term recording in rodents and nonhuman primates. Sci Transl Med, 12(538). https://doi.org/10.1126/scitranslmed.aay4682

Chung, J. E., Sellers, K. K., Leonard, M. K., Gwilliams, L., Xu, D., Dougherty, M. E., Kharazia, V., Metzger, S. L., Welkenhuysen, M., Dutta, B., & Chang, E. F. (2022). High-density single-unit human cortical recordings using the Neuropixels probe. Neuron, 110(15), 2409–2421. https://doi.org/10.1016/j.neuron.2022.05.007

Compte, A., Reig, R., Descalzo, V. F., Harvey, M. A., Puccini, G. D., & Sanchez-Vives, M. V. (2008). Spontaneous high-frequency (10-80 Hz) oscillations during up states in the cerebral cortex in vitro. J Neurosci, 28(51), 13828–13844. https://doi.org/10.1523/JNEUROSCI.2684-08.2008

Das, A., Myers, J., Mathura, R., Shofty, B., Metzger, B. A., Bijanki, K., Wu, C., Jacobs, J., & Sheth, S. A. (2022). Spontaneous neuronal oscillations in the human insula are hierarchically organized traveling waves. Elife, 11. https://doi.org/10.7554/eLife.76702

Fernandez, E., Alfaro, A., Soto-Sanchez, C., Gonzalez-Lopez, P., Lozano, A. M., Pena, S., Grima, M. D., Rodil, A., Gomez, B., Chen, X., Roelfsema, P. R., Rolston, J. D., Davis, T. S., & Normann, R. A. (2021). Visual percepts evoked with an intracortical 96-channel microelectrode array inserted in human occipital cortex. J Clin Invest, 131(23). https://doi.org/10.1172/JCI151331

Fernandez, E., Greger, B., House, P. A., Aranda, I., Botella, C., Albisua, J., Soto-Sanchez, C., Alfaro, A., & Normann, R. A. (2014). Acute human brain responses to intracortical microelectrode arrays: challenges and future prospects. Front Neuroeng, 7, 24. https://doi.org/10.3389/fneng.2014.00024

Flesher, S. N., Collinger, J. L., Foldes, S. T., Weiss, J. M., Downey, J. E., Tyler-Kabara, E. C., Bensmaia, S. J., Schwartz, A. B., Boninger, M. L., & Gaunt, R. A. (2016). Intracortical microstimulation of human somatosensory cortex. Sci Transl Med, 8(361), 361ra141. https://doi.org/10.1126/scitranslmed.aaf8083

Fries, P. (2015). Rhythms for Cognition: Communication through Coherence. Neuron, 88(1), 220–235. https://doi.org/10.1016/j.neuron.2015.09.034

Fu, Z., Beam, D., Chung, J. M., Reed, C. M., Mamelak, A. N., Adolphs, R., & Rutishauser, U. (2022). The geometry of domain-general performance monitoring in the human medial frontal cortex. Science, 376(6593), eabm9922. https://doi.org/10.1126/science.abm9922

Hill, D. N., Mehta, S. B., & Kleinfeld, D. (2011). Quality metrics to accompany spike sorting of extracellular signals. J Neurosci, 31(24), 8699–8705. https://doi.org/10.1523/JNEUROSCI.0971-11.2011

Hochberg, L. R., Serruya, M. D., Friehs, G. M., Mukand, J. A., Saleh, M., Caplan, A. H., Branner, A., Chen, D., Penn, R. D., & Donoghue, J. P. (2006). Neuronal ensemble control of prosthetic devices by a human with tetraplegia. Nature, 442(7099), 164–171. https://doi.org/10.1038/nature04970

House, P. A., MacDonald, J. D., Tresco, P. A., & Normann, R. A. (2006). Acute microelectrode array implantation into human neocortex: preliminary technique and histological considerations. Neurosurg Focus, 20(5), E4. https://doi.org/10.3171/foc.2006.20.5.5

Jacob, S. N., Hahnke, D., & Nieder, A. (2018). Structuring of Abstract Working Memory Content by Fronto-parietal Synchrony in Primate Cortex. Neuron, 99(3), 588–597. https://doi.org/10.1016/j.neuron.2018.07.025

Jacob, S. N., & Nieder, A. (2014). Complementary roles for primate frontal and parietal cortex in guarding working memory from distractor stimuli. Neuron, 83(1), 226–237. https://doi.org/10.1016/j.neuron.2014.05.009

Jamali, M., Grannan, B., Haroush, K., Moses, Z. B., Eskandar, E. N., Herrington, T., Patel, S., & Williams, Z. M. (2019). Dorsolateral prefrontal neurons mediate subjective decisions and their variation in humans. Nat Neurosci, 22(6), 1010–1020. https://doi.org/10.1038/s41593-019-0378-3

Jamali, M., Grannan, B. L., Fedorenko, E., Saxe, R., Baez-Mendoza, R., & Williams, Z. M. (2021). Single-neuronal predictions of others’ beliefs in humans. Nature, 591(7851), 610–614. https://doi.org/10.1038/s41586-021-03184-0

Kaminski, J., Sullivan, S., Chung, J. M., Ross, I. B., Mamelak, A. N., & Rutishauser, U. (2017). Persistently active neurons in human medial frontal and medial temporal lobe support working memory. Nat Neurosci, 20(4), 590–601. https://doi.org/10.1038/nn.4509

Kornblith, S., Quian Quiroga, R., Koch, C., Fried, I., & Mormann, F. (2017). Persistent Single-Neuron Activity during Working Memory in the Human Medial Temporal Lobe. Curr Biol, 27(7), 1026–1032. https://doi.org/10.1016/j.cub.2017.02.013

Kutter, E. F., Bostroem, J., Elger, C. E., Mormann, F., & Nieder, A. (2018). Single Neurons in the Human Brain Encode Numbers. Neuron, 100(3), 753–761. https://doi.org/10.1016/j.neuron.2018.08.036

Ledoit, O., & Wolf, M. (2004). A well-conditioned estimator for large-dimensional covariance matrices. Journal of Multivariate Analysis, 88(2), 365–411. https://doi.org/10.1016/s0047-259x(03)00096-4

Liou, J. Y., Smith, E. H., Bateman, L. M., McKhann, G. M., Goodman, R. R., Greger, B., Davis, T. S., Kellis, S. S., House, P. A., & Schevon, C. A. (2017). Multivariate regression methods for estimating velocity of ictal discharges from human microelectrode recordings. J Neural Eng, 14(4), 044001. https://doi.org/10.1088/1741-2552/aa68a6

Loomba, S., Straehle, J., Gangadharan, V., Heike, N., Khalifa, A., Motta, A., Ju, N., Sievers, M., Gempt, J., Meyer, H. S., & Helmstaedter, M. (2022). Connectomic comparison of mouse and human cortex. Science, 377(6602), eabo0924. https://doi.org/10.1126/science.abo0924

Mandonnet, E., & Herbet, G. (2021). Intraoperative Mapping of Cognitive Networks. Springer.

Meirhaeghe, N., Sohn, H., & Jazayeri, M. (2021). A precise and adaptive neural mechanism for predictive temporal processing in the frontal cortex. Neuron, 109(18), 2995–3011. https://doi.org/10.1016/j.neuron.2021.08.025

Minxha, J., Adolphs, R., Fusi, S., Mamelak, A. N., & Rutishauser, U. (2020). Flexible recruitment of memory-based choice representations by the human medial frontal cortex. Science, 368(6498), eaba3313. https://doi.org/10.1126/science.aba3313

Mitz, A. R., Bartolo, R., Saunders, R. C., Browning, P. G., Talbot, T., & Averbeck, B. B. (2017). High channel count single-unit recordings from nonhuman primate frontal cortex. J Neurosci Methods, 289, 39–47. https://doi.org/10.1016/j.jneumeth.2017.07.001

Moca, V. V., Barzan, H., Nagy-Dabacan, A., & Muresan, R. C. (2021). Time-frequency super-resolution with superlets. Nat Commun, 12(1), 337. https://doi.org/10.1038/s41467-020-20539-9

Muller, L., Chavane, F., Reynolds, J., & Sejnowski, T. J. (2018). Cortical travelling waves: mechanisms and computational principles. Nat Rev Neurosci, 19(5), 255–268. https://doi.org/10.1038/nrn.2018.20

Nieder, A. (2016). The neuronal code for number. Nat Rev Neurosci, 17(6), 366–382. https://doi.org/10.1038/nrn.2016.40

Nieder, A., Diester, I., & Tudusciuc, O. (2006). Temporal and spatial enumeration processes in the primate parietal cortex. Science, 313(5792), 1431–1435. https://doi.org/10.1126/science.1130308

Oostenveld, R., Fries, P., Maris, E., & Schoffelen, J. M. (2011). FieldTrip: Open source software for advanced analysis of MEG, EEG, and invasive electrophysiological data. Comput Intell Neurosci, 2011, 156869. https://doi.org/10.1155/2011/156869

Pandarinath, C., Nuyujukian, P., Blabe, C. H., Sorice, B. L., Saab, J., Willett, F. R., Hochberg, L. R., Shenoy, K. V., & Henderson, J. M. (2017). High performance communication by people with paralysis using an intracortical brain-computer interface. Elife, 6. https://doi.org/10.7554/eLife.18554

Paulk, A. C., Kfir, Y., Khanna, A. R., Mustroph, M. L., Trautmann, E. M., Soper, D. J., Stavisky, S. D., Welkenhuysen, M., Dutta, B., Shenoy, K. V., Hochberg, L. R., Richardson, R. M., Williams, Z. M., & Cash, S. S. (2022). Large-scale neural recordings with single neuron resolution using Neuropixels probes in human cortex. Nat Neurosci, 25(2), 252–263. https://doi.org/10.1038/s41593-021-00997-0

Piazza, M., Fumarola, A., Chinello, A., & Melcher, D. (2011). Subitizing reflects visuo-spatial object individuation capacity. Cognition, 121(1), 147–153. https://doi.org/10.1016/j.cognition.2011.05.007

Rubino, D., Robbins, K. A., & Hatsopoulos, N. G. (2006). Propagating waves mediate information transfer in the motor cortex. Nat Neurosci, 9(12), 1549–1557. https://doi.org/10.1038/nn1802

Rutishauser, U., Ross, I. B., Mamelak, A. N., & Schuman, E. M. (2010). Human memory strength is predicted by theta-frequency phase-locking of single neurons. Nature, 464(7290), 903–907. https://doi.org/10.1038/nature08860

Sanai, N., Mirzadeh, Z., & Berger, M. S. (2008). Functional outcome after language mapping for glioma resection. N Engl J Med, 358(1), 18–27. https://doi.org/10.1056/NEJMoa067819

Sato, T. K., Nauhaus, I., & Carandini, M. (2012). Traveling waves in visual cortex. Neuron, 75(2), 218–229. https://doi.org/10.1016/j.neuron.2012.06.029

Schevon, C. A., Tobochnik, S., Eissa, T., Merricks, E., Gill, B., Parrish, R. R., Bateman, L. M., McKhann, G. M., Jr., Emerson, R. G., & Trevelyan, A. J. (2019). Multiscale recordings reveal the dynamic spatial structure of human seizures. Neurobiol Dis, 127, 303–311. https://doi.org/10.1016/j.nbd.2019.03.015

Shattuck, D. W., & Leahy, R. M. (2002). BrainSuite: An automated cortical surface identification tool. Medical Image Analysis, 6(2), 129–142. https://doi.org/10.1016/s1361-8415(02)00054-3

Sheth, S. A., Mian, M. K., Patel, S. R., Asaad, W. F., Williams, Z. M., Dougherty, D. D., Bush, G., & Eskandar, E. N. (2012). Human dorsal anterior cingulate cortex neurons mediate ongoing behavioural adaptation. Nature, 488(7410), 218–221. https://doi.org/10.1038/nature11239

Smith, E. H., Liou, J. Y., Davis, T. S., Merricks, E. M., Kellis, S. S., Weiss, S. A., Greger, B., House, P. A., McKhann, G. M., 2nd, Goodman, R. R., Emerson, R. G., Bateman, L. M., Trevelyan, A. J., & Schevon, C. A. (2016). The ictal wavefront is the spatiotemporal source of discharges during spontaneous human seizures. Nat Commun, 7, 11098. https://doi.org/10.1038/ncomms11098

Suarez-Perez, A., Gabriel, G., Rebollo, B., Illa, X., Guimera-Brunet, A., Hernandez-Ferrer, J., Martinez, M. T., Villa, R., & Sanchez-Vives, M. V. (2018). Quantification of Signal-to-Noise Ratio in Cerebral Cortex Recordings Using Flexible MEAs With Co-localized Platinum Black, Carbon Nanotubes, and Gold Electrodes. Front Neurosci, 12, 862. https://doi.org/10.3389/fnins.2018.00862

Takahashi, K., Saleh, M., Penn, R. D., & Hatsopoulos, N. G. (2011). Propagating waves in human motor cortex. Front Hum Neurosci, 5, 40. https://doi.org/10.3389/fnhum.2011.00040

Tong, A. P. S., Vaz, A. P., Wittig, J. H., Inati, S. K., & Zaghloul, K. A. (2021). Ripples reflect a spectrum of synchronous spiking activity in human anterior temporal lobe. Elife, 10. https://doi.org/10.7554/eLife.68401

Truccolo, W., Donoghue, J. A., Hochberg, L. R., Eskandar, E. N., Madsen, J. R., Anderson, W. S., Brown, E. N., Halgren, E., & Cash, S. S. (2011). Single-neuron dynamics in human focal epilepsy. Nat Neurosci, 14(5), 635–641. https://doi.org/10.1038/nn.2782

Vaz, A. P., Wittig, J. H., Jr., Inati, S. K., & Zaghloul, K. A. (2020). Replay of cortical spiking sequences during human memory retrieval. Science, 367(6482), 1131–1134. https://doi.org/10.1126/science.aba0672

Willett, F. R., Avansino, D. T., Hochberg, L. R., Henderson, J. M., & Shenoy, K. V. (2021). High-performance brain-to-text communication via handwriting. Nature, 593(7858), 249–254. https://doi.org/10.1038/s41586-021-03506-2

Woods, B. (2011). Spatio-temporal Patterns in Multi-Electrode Array Local Field Potential Recordings. arXiv, 1501.00230v1.

Zaghloul, K. A., Blanco, J. A., Weidemann, C. T., McGill, K., Jaggi, J. L., Baltuch, G. H., & Kahana, M. J. (2009). Human substantia nigra neurons encode unexpected financial rewards. Science, 323(5920), 1496–1499. https://doi.org/10.1126/science.1167342

Zaghloul, K. A., Weidemann, C. T., Lega, B. C., Jaggi, J. L., Baltuch, G. H., & Kahana, M. J. (2012). Neuronal activity in the human subthalamic nucleus encodes decision conflict during action selection. J Neurosci, 32(7), 2453–2460. https://doi.org/10.1523/JNEUROSCI.5815-11.2012

Zhang, H., & Jacobs, J. (2015). Traveling Theta Waves in the Human Hippocampus. J Neurosci, 35(36), 12477–12487. https://doi.org/10.1523/JNEUROSCI.5102-14.2015

Zhang, H., Watrous, A. J., Patel, A., & Jacobs, J. (2018). Theta and Alpha Oscillations Are Traveling Waves in the Human Neocortex. Neuron, 98(6), 1269–1281. https://doi.org/10.1016/j.neuron.2018.05.019

Zilio, F., Gomez-Pilar, J., Cao, S., Zhang, J., Zang, D., Qi, Z., Tan, J., Hiromi, T., Wu, X., Fogel, S., Huang, Z., Hohmann, M. R., Fomina, T., Synofzik, M., Grosse-Wentrup, M., Owen, A. M., & Northoff, G. (2021). Are intrinsic neural timescales related to sensory processing? Evidence from abnormal behavioral states. Neuroimage, 226, 117579. https://doi.org/10.1016/j.neuroimage.2020.117579

